# Fibroblasts generate topographical cues that steer cancer cell migration

**DOI:** 10.1101/2022.09.06.506801

**Authors:** Francesco Baschieri, Abigail Illand, Jorge Barbazan, Olivier Zajac, Clémence Henon, Damarys Loew, Florent Dingli, Danijela Matic Vignjevic, Sandrine Lévêque-Fort, Guillaume Montagnac

## Abstract

Fibroblasts play a fundamental role in tumor development. Among other functions, they regulate cancer cells migration through rearranging the extracellular matrix, secreting soluble factors and establishing direct physical contacts with cancer cells. Here, we report that migrating fibroblasts deposit on the substrate a network of tubular structures that serves as guidance cue for cancer cell migration. Such membranous tubular network, hereafter called tracks, is stably anchored to the substrate in a β5 integrin-dependent manner. We found that cancer cells specifically adhere to tracks by using clathrin-coated structures that pinch and engulf tracks. Tracks represent thus a spatial memory of fibroblast migration paths that is read and erased by cancer cells directionally migrating along them. We propose that fibroblast tracks represent a topography-based intercellular communication system capable of steering cancer cells migration.

**TEASER:** The migration path of fibroblasts is marked by tubules that act as railways to direct following cancer cell migration.

## MAIN TEXT

Cell migration is fundamental in cancer development as it is linked to tumor cell invasion and formation of metastases (Fares et al., 2020). Several chemical and physical cues of the tumor microenvironment were described to regulate cancer cell migration (Erdogan et al., 2017; Fischer et al., 2021; Oudin & Weaver, 2016; Wyckoff et al., 2004). Fibroblasts, and in particular cancer-associated fibroblasts (CAFs), act on cancer cell migration in several ways, sometimes preventing invasion by building a capsule of extracellular matrix around the tumor (Zhang et al., 2013), and sometimes favoring cancer cells escape from the primary tumor by secreting soluble factors (Erdogan et al., 2017), modifying the extracellular matrix (Attieh et al., 2017; Gaggioli et al., 2007) or physically pulling on cancer cells (Labernadie et al., 2017). In this latter case, CAFs migration directly influences cancer cells migration. Fibroblasts were shown to deposit a membranous material composed of tubules of plasma membrane emanating from retraction fibers at the cell rear during migration (Fuhr et al., 1998; Zimmermann et al., 2001). More recently, coalescence of these tubules into large vesicles termed migrasomes were described in migrating fibroblasts (Huang et al., 2019; Jiang et al., 2019; Ma et al., 2015). Migrasomes are short-lived structures that rupture soon after their formation and release their content of signaling molecules to orientate the migration of following cells (Jiang et al., 2019). Migrasomes can also be internalized by surrounding cells and thereby transfer regulatory factors they contain to modulate recipient cells fate (Jiang et al., 2019; Yu & Yu, 2021). While it is proven that fibroblasts can produce migrasomes, it is not clear whether these structures may participate in orienting cancer cell migration. In addition, depending on the cell type, migrasomes are often less numerous than the total amount of membranous tubules left behind by fibroblasts (Fan et al., 2022). This opens the possibility that the tubular part of the network also plays a role in directing following cells. Here, we set out to determine if and how membranous materials left on the substrate by migrating CAFs can steer cancer cell migration.

## RESULTS

We observed that immortalized CAFs isolated from a colon cancer patient left behind themselves an extended network of membranous material while migrating on glass (Movie 1 and Figure 1A). The migration path of CAFs evolving in a 3D network of collagen fibers was also decorated by similar membranous structures, indicating that these structures do not only form in artificial 2D setups (Supp. Fig. 1A). This network originated from retraction fibers as the cell moved forwards (Movie 1 and Figure 1A). Pearlation of tubules composing the network was occasionally observed suggesting that tubules are under tension (Bar-Ziv & Moses, 1994) (Supp. Fig. 1B). The network was organized as branches of regular angles pointing in the direction of the CAF that left them on the substrate (Figure 1B and C). Confirming previous findings (Zimmermann et al., 2001), we measured by super resolution microscopy that tubules composing this network displayed a very regular width (Supp. Fig 1C and D). We also noticed some rare and large vesicles seemingly connected to the branches (Figure 1C). However, in agreement with the literature (Ma et al., 2015), we observed that these vesicles, most likely corresponding to migrasomes, quickly disappeared after their formation while the branched network remained stable for days (Figure 1C-E). We hereafter refer to this remaining network as tracks. Filamentous actin and microtubules were not present in tracks (Supp. Fig. 2A and B). Interference reflection microscopy revealed that tracks deposited on glass by CAFs established very tight contacts with the substrate suggesting that they specifically adhere to it (Supp. Fig. 2C). Thus, we stained CAFs and tracks for different integrins and observed that β5 integrin but not β1 nor β3 integrins accumulated in tracks (Figure 1F). We next observed that integrin β5 knockdown completely abrogated track biogenesis (Figure 1G and H and Supp. Fig. 2D and E). Conversely, β1 depletion or knock-out did not prevent track formation by fibroblasts (Supp. Fig. 2F-I). Of note, β5-integrin -positive tracks were produced by CAFs from different origins, as well as by mesenchymal stem cells and immortalized mouse osteoblasts (Supp. Fig. 2J). Thus, our results suggest that tracks are adhesive membranous networks anchored to the substrate in an integrin β5-dependent manner.

**FIGURE 1.**
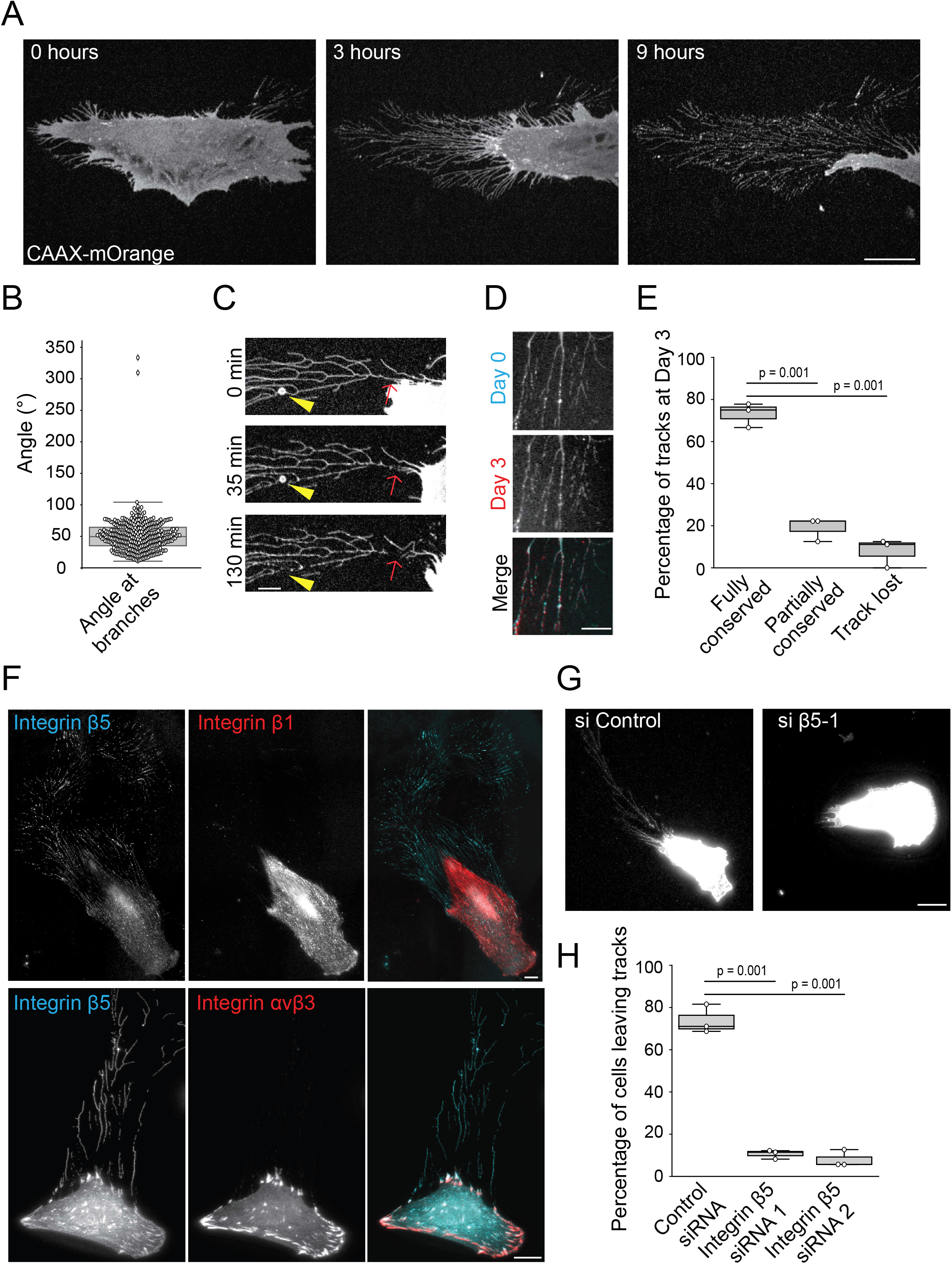
CAFs deposit tracks in a β5-integrin-dependent manner. **A**, CAF stably expressing CAAX-mOrange were seeded on glass and imaged by spinning disk microscopy for 9 hours. Scale bar: 20 μm. **B**, Measurement of angles formed by track-branches. **C**, CAF stably expressing CAAX-mOrange were allowed to migrate on glass and imaged by spinning disk microscopy. Arrowheads point to a migrasome and arrows indicate the site of membrane rupture. Scale bar: 5μm. **D**, CAFs stably expressing BFP-tagged β5-integrin were allowed to deposit tracks on glass. Tracks were imaged the first day (Day 0) and then at day 3 after deposition. Scale bar: 10 μm. **E**, Tracks as in D were imaged on Day 0 and Day 3. The histogram represents the percentage of tracks being identical between the two time points (Fully conserved), deteriorated (Partially conserved) or lost. A total of 26 tracks from 3 independent experiments were monitored. The percentage of each category is shown on the graph ± SD (Kruskal-Wallis test). **F**, CAFs were allowed to migrate on glass, then fixed and stained with the indicated antibodies. Scale bars: 20 μm. **G**. CAFs transfected with control or β5-integrin siRNA were allowed to migrate on glass for 48h, before cells and tracks were labelled with Alexa-488-labeleld Wheat Germ Agglutinin and imaged. Scale bar: 20 μm. **H**. Cells as in G were monitored for track deposition. Data represent the mean percentage ± SD of cells depositing tracks from three independent experiments (Kruskal-Wallis test).

We next investigated how cancer cells behave upon contacting CAF-tracks. First, we tested a possible role for tracks serving as adhesion cues for cancer cells. For this, CAFs were allowed to deposit tracks on the substrate for 24h. CAFs were then detached using a procedure that left tracks intact on the substrate (see Material and Method section). MDA-MB-231 cancer cells were then seeded on the CAF-conditioned substrate and imaged to monitor sites of cell spreading relative to the position of tracks. MDA-MB-231 cells showed a strong preference for adhering in track regions as compared to other areas of the substrate (Figure 2A and B). We also noticed that upon adhering and spreading on tracks, MDA-MB-231 cells migrated by following the tracks (Figure 2C and Movie 2).

**FIGURE 2.**
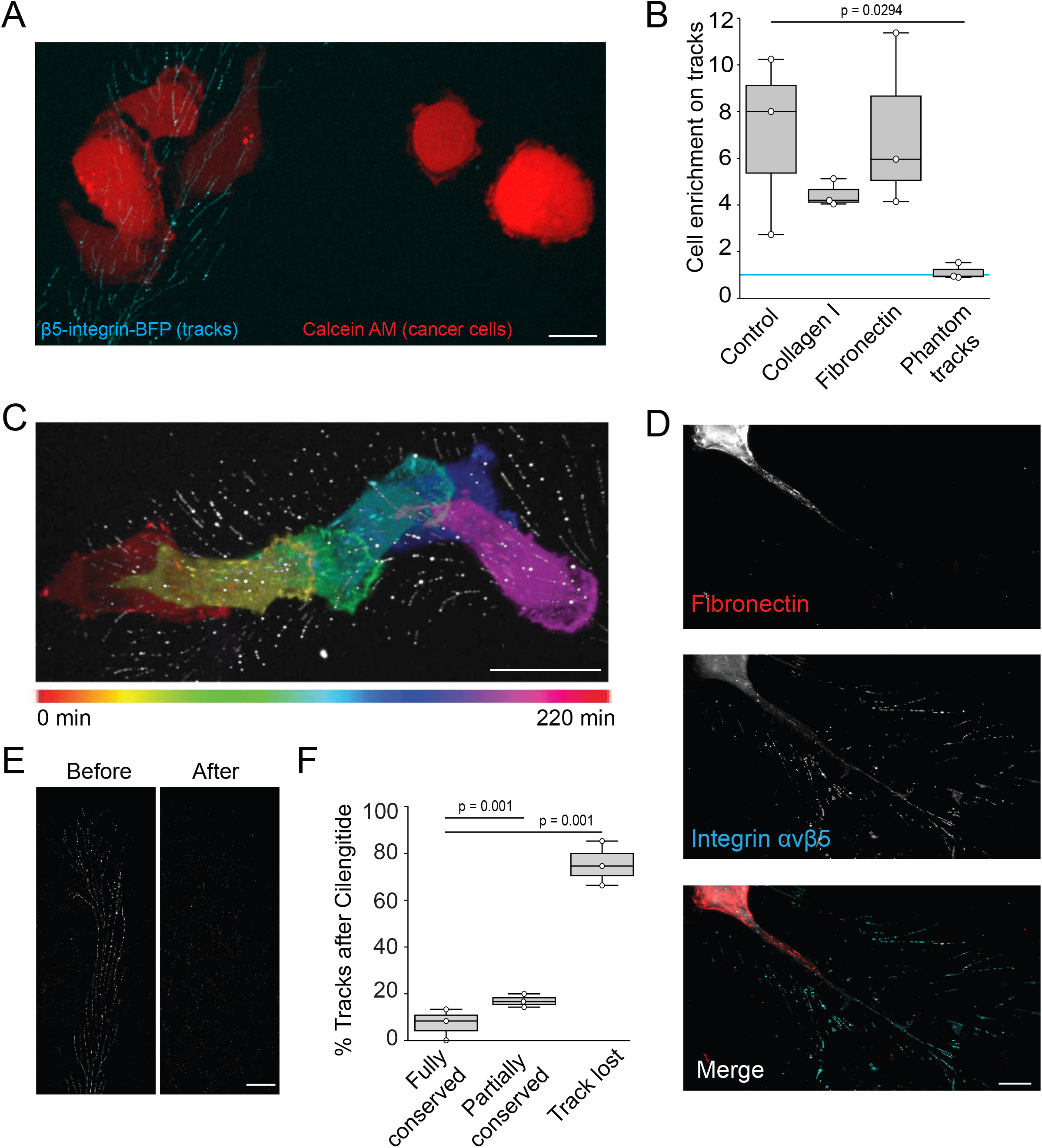
CAF-tracks represent adhesion cues for cancer cells. **A**, MDA-MB-231 cells marked with Calcein AM were allowed to spread on glass substrates covered with tracks deposited by β5-integrin-BFP overexpressing CAFs. Scale bar: 10 μm. **B**, β5-integrin-BFP overexpressing CAFs were allowed to deposit tracks for 48 h on glass-bottom dishes coated or not with collagen or fibronectin, as indicated. Tracks positions were imaged before MDA-MB-231 cells were seeded on the substrates. For the “Phantom tracks” condition, tracks were imaged and removed from the substrate using Cilengitide before seeding MDA-MB-231 cells. The number of MDA-MB-231 cells adhering in tracks areas versus in other areas of the substrates was measured. Results are expressed as the mean ratio ± SD of cell density on tracks versus in other areas of the substrate from three independent experiments (Uncorrected Fisher’s LSD test). A value of 1 (blue line) means no enrichment on tracks. **C**, Color-coded representation of time-lapse recording of vinculin-GFP-expressing MDA-MB-231 cells allowed to migrate for 220 min on tracks deposited by β5-integrin-BFP expressing CAFs. Scale bar: 20 μm. **D**, CAFs were allowed to deposit tracks on glass before to be stained for αvβ5 integrin and fibronectin, as indicated. Scale bar: 10 μm. **E**, CAFs expressing β5-integrin-BFP were allowed to migrate for 48h on gridded coverslips. Tracks positions were recorded before Cilengitide was added to remove tracks and same positions were imaged again (after). Scale bar: 20 μm. **F**, Tracks as in E and originating from 41 cells from 3 independent experiments were evaluated for their integrity after Cilengitide treatment. The percentage of each category ± SD is shown (Kruskal-Wallis test).

In addition to tracks, CAFs are known to deposit extracellular matrix components that may affect cancer cell adhesion (Attieh et al., 2017; Erdogan et al., 2017). However, we noticed that fibronectin was virtually not deposited by CAFs in the low confluency conditions used to visualize tracks (Figure 2D). To exclude the possibility that cancer cells were sensing undetected extracellular matrix deposits, we first allowed CAFs to deposit tracks on collagen-or fibronectin-coated glass-coverslips. The matrix coating slightly reduced the preferential adhesion of cancer cells to tracks regions although the differences were not statistically significant (Figure 2B). This suggests that tracks represent more effective adhesion cues than collagen or fibronectin. To ensure that tracks were sufficient to support adhesion in tracks areas, we next recorded the position of tracks deposited by CAFs before tracks were removed using Cilengitide. Indeed, Cilengitide, a competitive inhibitor of αvβ5 integrin, allowed to specifically detach tracks from the substrate (Figure 2E and F). We observed that MDA-MB-231 cells did not preferentially spread in regions previously occupied by tracks (Figure 2B). This demonstrates that tracks are required and sufficient for the observed preferential adhesion in tracks areas. Together, our results show that tracks are very potent adhesion cues for cancer cells.

We next aimed to determine how cancer cells adhere to tracks. We first monitored the distribution of focal adhesions marked with vinculin or Talin1 in MDA-MB-231 cells adhering on tracks. Although focal adhesions were present in these cells, they never overlapped with tracks (Figure 3A and B and Supp. Fig. 3A and B). Additionally, the ability of cells to preferentially adhere to tracks remained unchanged upon depletion of Talin1, an essential component of focal adhesions (Figure 3C and Supp. Fig. 3C). These results suggest that focal adhesions do not regulate cancer cells adhesion on tracks. We recently showed that clathrin-coated structures (CCSs) accumulate along collagen fibers where they serve as adhesion structures in an integrin β1-dependent manner (Elkhatib et al., 2017). Because tracks are tubular structures and show a similar diameter to collagen fibers (Supp. Fig. 1C), we hypothesized that CCSs of cancer cells could also serve as adhesion structures to tracks. First, we observed that CCSs of MDA-MB-231 cells marked with the μ2-adaptin subunit of the AP-2 complex strongly accumulated along tracks (Figure 3D and E). Similar CCSs accumulation were observed in HCT116 colon cancer cells adhering on tracks (Supp. Fig. 3D and E) as well as in MDA-MB-231 adhering on tracks deposited by osteoblasts, another type of fibroblastic cells (Supp. Fig. 3F and G). We observed that both CCSs lifetime and nucleation rate were increased at cancer cell/tracks contact sites as compared to other areas of the plasma membrane (Figure 3F and G and Supp. Fig. 3H and I). This phenotype is remarkably similar to collagen fibers-pinching CCSs and suggests that CCSs could play a role in cancer cell adhesion to tracks. Indeed, CCSs ablation by knock-down of the CCSs core component AP-2 strongly reduced the preferential adhesion of MDA-MB-231 cells to tracks (Figure 3C). However, depletion of clathrin heavy chain (CHC) did not inhibit preferential adhesion to tracks (Figure 3C). We already reported that CHC, although required for CCSs budding and endocytosis, is not required for the formation of AP-2 lattices that can serve as adhesion structures to collagen fibers (Bresteau et al., 2021; Elkhatib et al., 2017). Our results suggest that CCSs on tracks, similarly to CCSs on collagen fibers, support adhesion to tracks in an endocytosis-independent manner.

**FIGURE 3.**
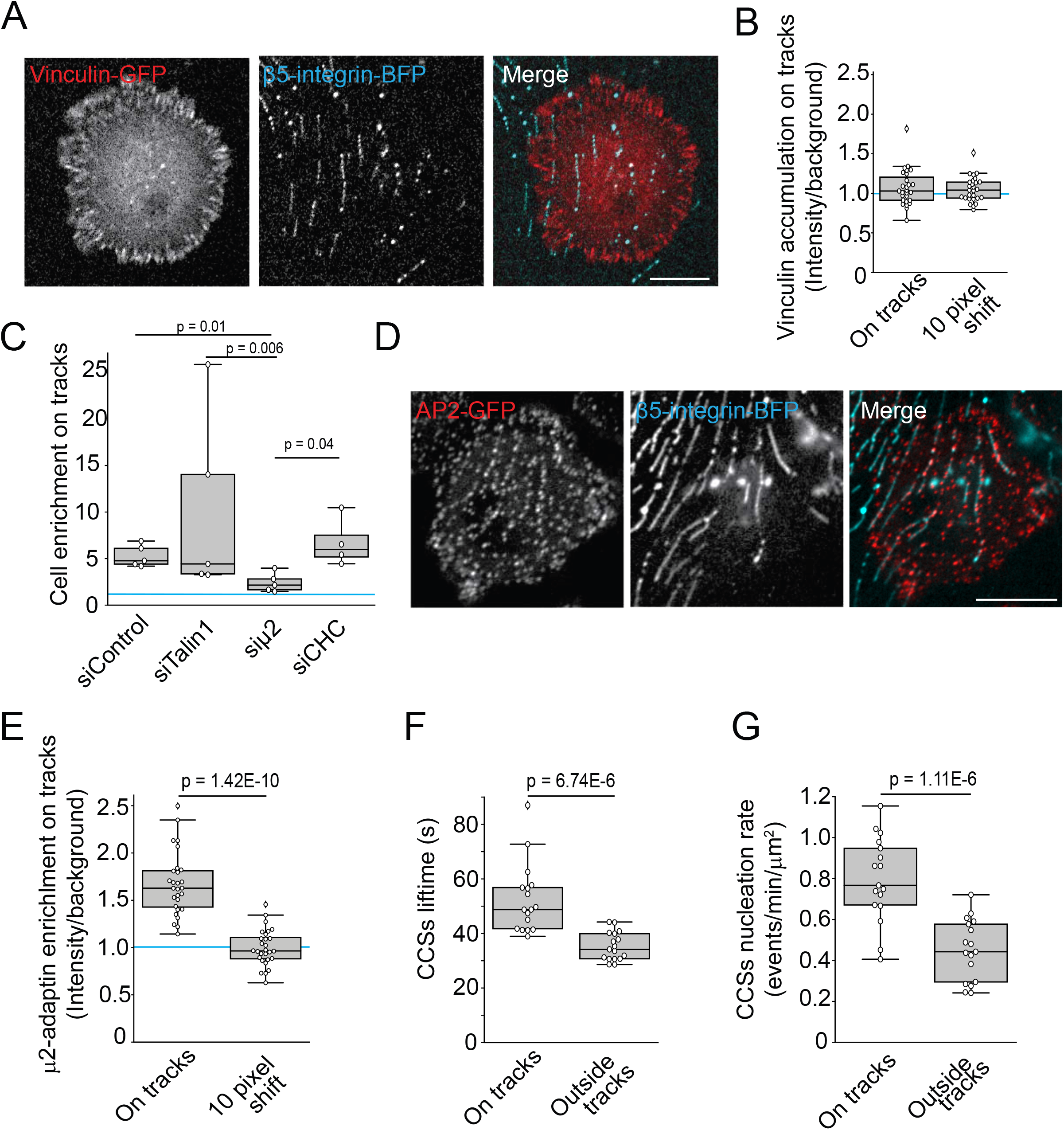
CCSs of MDA-MB-231 cells accumulate on and are required to adhere to CAF-tracks. **A**, Vinculin-GFP expressing MDA-MB-231 cells were allowed to spread on β5-integrin-BFP-positive tracks. Images were acquired by spinning disk microscopy 30 min after seeding cells. Scale bar: 10 μm **B**, Quantification of vinculin-associated fluorescence in track areas versus other areas of the plasma membrane in cells as in A. Quantifications were performed with aligned vinculin and β5-integrin signals (on tracks) as well as upon shifting the β5-integrin signal by 10 pixels (10 pixels shift). 27 cells from three independent experiments were quantified. A value of 1 (blue line) means no enrichment on tracks. Results are represented as mean ratio of track-associated versus non-track-associated vinculin signal ± SD (Student’s T-test). **C**, MDA-MB-231 cells transfected with the indicated siRNAs were seeded on β5-integrin-BFP and imaged every 5 min for 8 hours. Results are expressed as mean ratio of cell number in tracks versus non-tracks areas ± SD from five independent experiments (Kruskal-Wallis test). A value of 1 (blue line) means no enrichment on tracks. **D**, MDA-MB-231 genome-edited to express μ2-adaptin-mCherry were allowed to spread on β5-integrin-BFP tracks and imaged 30 min after seeding. Scale bar: 10 μm. **E**, Quantification of μ2-adaptin-associated fluorescence in track areas versus other areas of the plasma membrane in cells as in D. Quantifications were performed with aligned μ2-adaptin and β5-integrin signals (on tracks) as well as upon shifting the β5-integrin signal by 10 pixels (10 pixels shift). 28 cells from 3 independent experiments were analyzed. A value of 1 (blue line) means no enrichment on tracks. Results are represented as mean ratio of track-associated versus non-track-associated μ2-adaptin signal ± SD (Student’s T-test). **F, G**, MDA-MB-231 genome-edited to express μ2-adaptin-mCherry were allowed to spread on β5-integrin-BFP tracks and imaged every 5 sec for 5 min. Lifetime (F) and nucleation rates (G) of CCSs on tracks vs outside of tracks were measured. 17 cells from three independent experiments were analyzed. Results are expressed as mean ± SD (Student’s T-test).

We next observed by super resolution microscopy that CCSs wrapped around and pinched tracks labelled with β5-integrin (Figure 4A and movie 3). We also noticed that some CCSs forming on tracks showed a budded profile and were filled with integrin β5 (Figure 4B), suggesting that CCSs can internalize portions of tracks. Indeed, we observed that tracks progressively disappeared below MDA-MB-231 cells adhering on them and this was dependent on both AP-2 and clathrin expression (Figure 4C). Our results suggest that CCSs on tracks experience a transient period of frustration, as reflected by their longer lifetime, but eventually manage to internalize portions of tracks. Actin has been proposed to be recruited at frustrated CCSs that experience difficulties to bud because of mechanical constraints (Baschieri et al., 2020; Boulant et al., 2011; Jin et al., 2022; Kaksonen et al., 2006; Perrais & Merrifield, 2005). Indeed, we observed that inhibiting actin dynamics with Cytochalasin D resulted in an increased accumulation of stable CCSs along tracks (Supp. Fig. 4A and B). In addition, Cytochalasin D incubation prevented tracks disappearance below MDA-MB-231 cells (Supp. Fig. 4C and movie 4). Thus, our results suggest that actin dynamics assists CCSs budding on tracks and allows tracks uptake.

**FIGURE 4.**
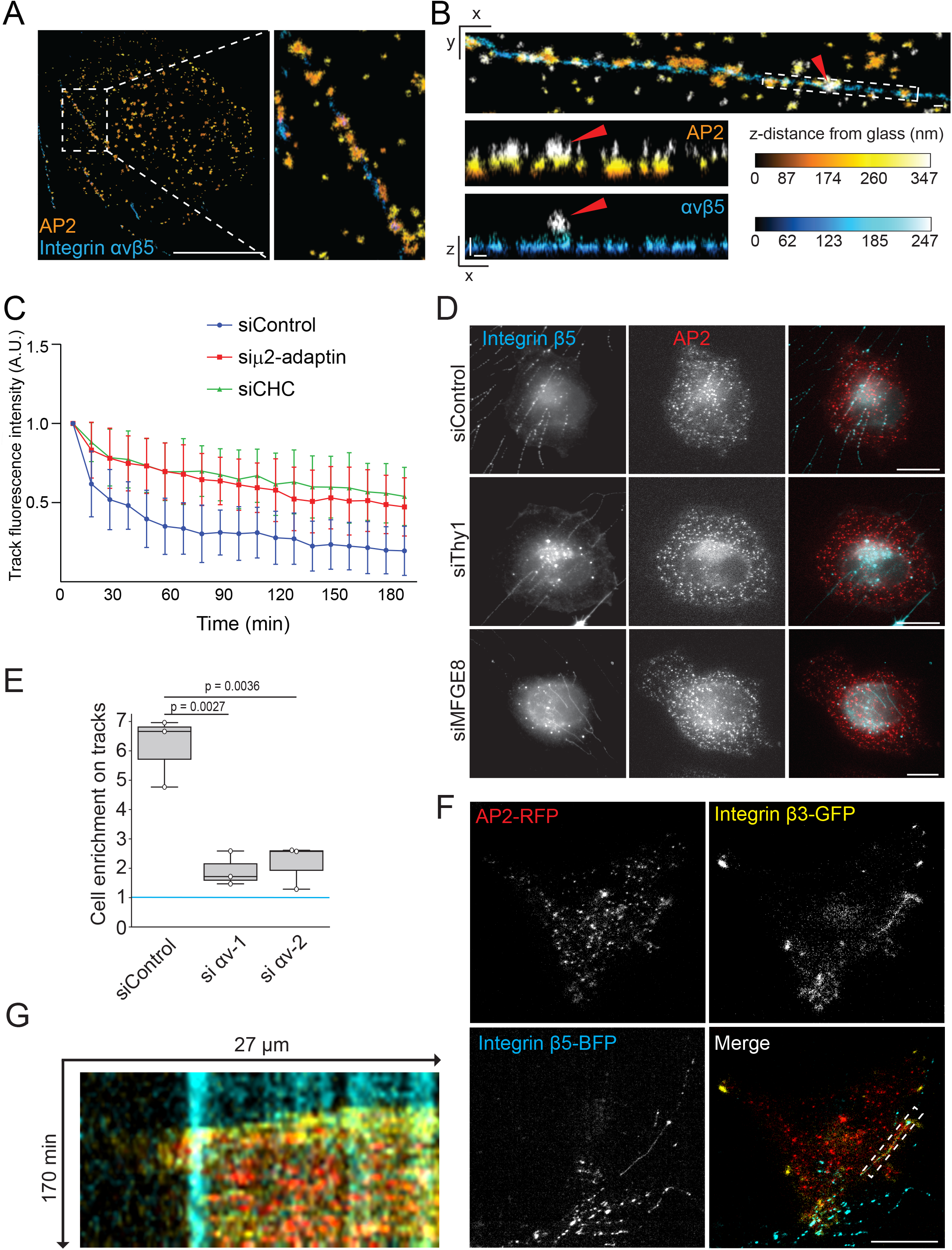
Mechanism of CCSs recruitment and cancer cell adhesion on tracks. **A**, Immunostaining of the α-adaptin subunit of AP-2 (MDA-MB-231 cells) and integrin αvβ5 (CAF-track) imaged by 3D super-resolution microscopy. Scale bar: 10 μm. **B**, 3D super-resolution microscopy images of α-adaptin and integrin αvβ5 as in A and observed in XY and XZ planes, as indicated. Red arrowheads point to a budding CCS containing integrin αvβ5. XZ projections of the boxed area are shown. Scale bars: 200 nm. Color-coded scales indicate the distance from glass in nm. **C**, MDA-MB-231 transfected with the indicated siRNAs were allowed to spread on β5-integrin-BFP tracks and imaged every 10 min for 3 hours. Data represent the evolution over time of the mean β5-integrin-BFP fluorescence intensity ± SD from three independent experiments (Dunnett’s multiple comparisons test; P<0.0001 as compared to siControl). **D**, CAFs transfected with the indicated siRNAs were allowed to migrate on glass coverslips for 48 h to deposit tracks. MDA-MB-231 genome-edited to express μ2-adaptin-GFP-RFP were then seeded on tracks for 35 min before to be fixed and tracks were stained with β5-integrin specific antibodies. Scale bars: 10 μm. **E**, MDA-MB-231 transfected with the indicated siRNAs were allowed to spread on β5-integrin-BFP tracks and imaged every 5 min for 8 hours. The number of cancer cells adhering on tracks versus outside of tracks was quantified and expressed as a mean ratio ± SD from three independent experiments (Kruskal-Wallis test). A value of 1 (blue line) means no enrichment on tracks. **F**, MDA-MB-231 genome-edited to express μ2-adaptin-mCherry (red) and stably expressing β3-integrin-GFP (yellow) were allowed to spread on β5-integrin-BFP osteoblast-tracks (cyan) and imaged by TIRF microscopy every 5 min for 8 hours. Representative images of the initial phase of MDA-MB-231 cells spreading on tracks are shown. Scale bar: 20 μm. **G**, Kymograph corresponding to the boxed area in F.

We next decided to investigate how CCSs are recruited on tracks. We reasoned that some CCS cargos may engage proteins expressed at the surface of tracks. We thus performed mass spectrometry analyses of purified tracks (see Material and Methods) to identify potential candidates. First, we observed that the fraction enriched for tracks contained integrin β5 and its partner integrin αv (Supp. Table 1), which was then confirmed by western blot analyses (Supp. Fig. 4D). Among the identified hits, we focused on plasma membrane-associated proteins involved in cell-cell adhesion. CAFs depleted for these proteins could still deposit tracks, so we seeded cancer cells on these tracks and measured CCSs accumulation on tracks. We observed a reduced accumulation of CCSs along tracks depleted of Thy1 or MFGE8 proteins (Figure 4D and Supp. Fig. 4E). Expression of GFP-tagged Thy1 and MFGE8 in CAFs confirmed that these proteins were present in tracks (Supp. Fig. 4F). Both Thy1 and MFGE8 were reported to bind in *trans* to the αvβ3 integrin dimer (Hanayama et al., 2002; Hermosilla et al., 2008; Leyton et al., 2001). We first confirmed that both MDA-MB-231 and HCT116 expressed αv and β3 integrins (Supp. Fig. 5G). We next observed that integrin αv knockdown reduced the preferential adhesion of MDA-MB-231 cells on wild-type tracks (Figure 4E and Supp. Fig. 4H). In addition, we observed that GFP-tagged integrin β3 strongly but transiently accumulated along tracks during cancer cell spreading (Movie 5). Occasional colocalization between CCSs and GFP-β3 was observed during MDA-MB-231 cells spreading on tracks (Figure 4F and G and movie 5). In addition, inhibiting tracks uptake with Cytochalasin D led to an increased and sustained accumulation of GFP-β3 along tracks (Supp. Fig. 4I and J). These latter observations suggest that αvβ3 transiently accumulate along tracks before to be removed through endocytosis, most likely together with portions of tracks. Overall, our results show that αvβ3 in MDA-MB-231 cells and αvβ3-interacting receptors on tracks are required for CCSs-dependent cancer cell adhesion to tracks.

Our findings suggest that tracks represent adhesion cues for cancer cells. We reasoned that such adhesion cues may also orient cancer cell migration. Fibroblasts perform durotaxis when seeded on substrates presenting non-homogenous rigidities (DuChez et al., 2019; Lo et al., 2000; Plotnikov et al., 2012). We first confirmed that CAFs efficiently migrated towards the stiffer end of a rigidity gradient (Figure 5A and Supp. Fig. 5A and B). However, MDA-MB-231 and HCT116 cells did not durotax on the rigidity gradient but rather migrated randomly (Figure 5A). We also observed that CAFs deposited more tracks on stiff than on soft substrates (Figure 5B). To determine whether tracks can orient cancer cells migration, CAFs were allowed to migrate and deposit track on the rigidity gradient before to be removed. We then seeded cancer cells on the CAF-conditioned rigidity gradient and monitored their migration. We observed that both MDA-MB-231 and HCT116 durotaxed on the CAF-conditioned gradient (Figure 5C and D). Similar results were obtained when MDA-MB-231 cells migrated on an osteoblast-conditioned rigidity gradient (Supp. Fig. 5C and Fig. 5A). However, Cilengitide-mediated tracks removal before cancer cells seeding resulted in both cell lines migrating randomly (Figure 5C and D). Thus, we concluded that tracks deposited by CAFs regulate the direction of cancer cells migration. We next tested the role of CCSs in this directed migration assay. We observed that AP-2 subunit μ2-adaptin knockdown prevented MDA-MB-231 cells to durotax on the CAF-conditioned gradient (Figure 5C). However, in line with the non-requirement for clathrin in mediating adhesion to tracks (Figure 3C), CHC-depleted cells still performed durotaxis on the CAF-conditioned gradient (Figure 5C). Collectively, our results show that tracks guide the migration of cancer cells and that this is regulated by CCS-mediated adhesion to tracks.

**FIGURE 5.**
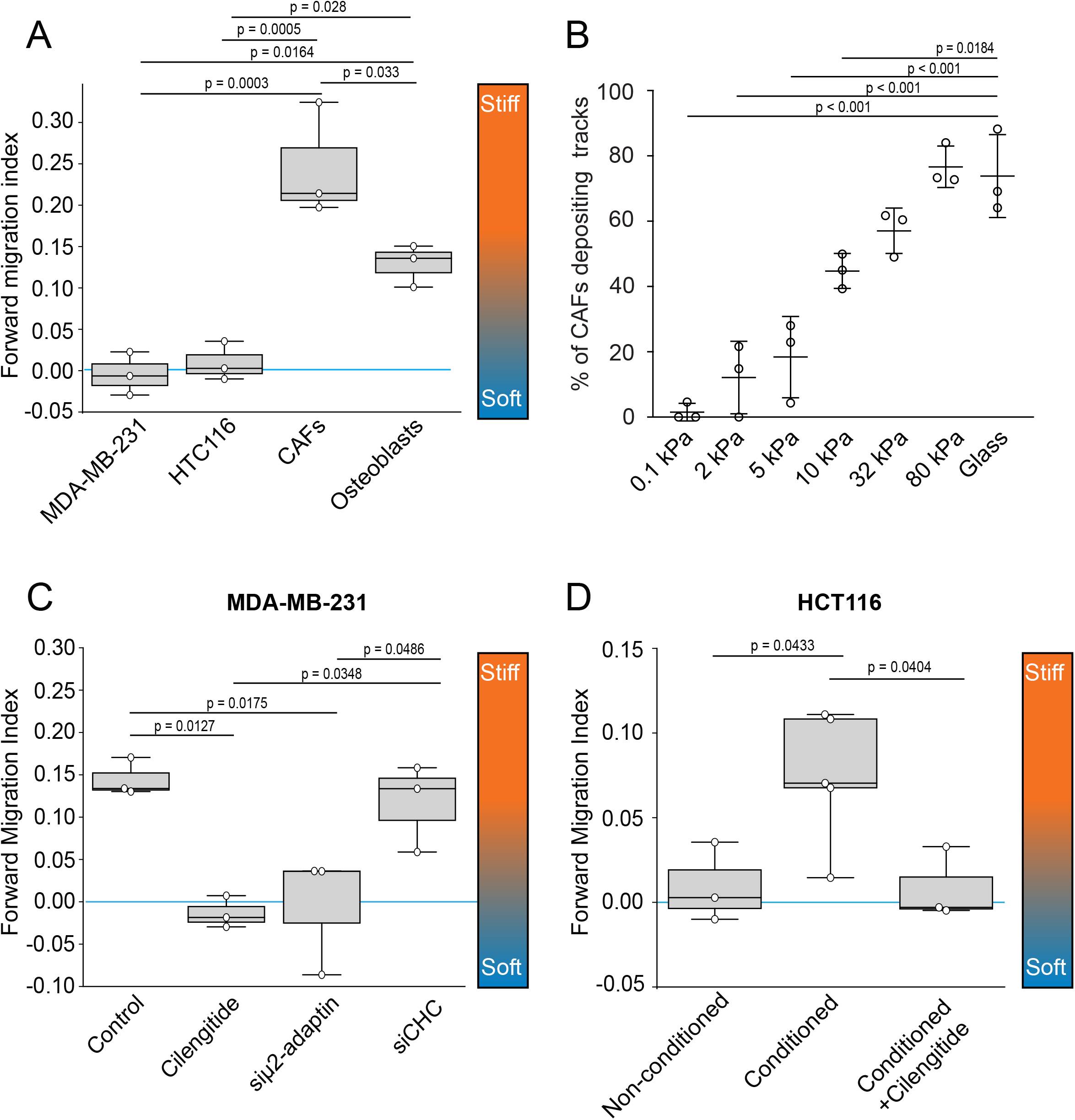
CAF-Tracks orientate cancer cell migration. **A**, Indicated cell lines were allowed to migrate on a collagen-coated rigidity gradient (0.1-80 kPa) and imaged every 20 min for 14 hours (see Materials and Methods section). Cells were manually tracked and the forward migration index (FMI) was calculated using the Chemotaxis tool plugin in FIJI software. Values close to 0 (blue line) indicate a random migration. Orientation of the gradient is shown. A total of 350 MDA-MB-231 cells, 204 CAFs, 514 HTC116 cells, and 505 osteoblasts were tracked in three independent experiments. Data are expressed as mean FMI ± SD (ANOVA one way, Turkey’s multiple comparison). **B**, CAFs were plated on collagen-coated polyacrylamide gels of indicated rigidities and allowed to deposit tracks for 24 hours before to be fixed and stained for β5-integrin. Results are expressed as the mean percentage of cells depositing tracks ± SD from three independent experiments (ANOVA one way, Turkey’s multiple comparison). **C**, Control or siRNA-treated MDA-MB-231 cells, as indicated, were allowed to migrate on CAF-conditioned rigidity gradients and imaged every 20 min for 14 hours. In the Cilengitide condition, deposited tracks were removed with Cilengitide before MDA-MB-231 were seeded on the gradient. Cells were manually tracked and forward migration index (FMI) was calculated using the Chemotaxis tool plugin of FIJI. Values close to 0 (blue line) indicate a random migration. Orientation of the gradient is shown. 281, 626, 424, and 362 cells were tracked from three independent experiments in the control, Cilengitide, siμ2-adaptin and siCHC conditions, respectively. Data are expressed as mean FMI ± SD (ANOVA one way, Turkey’s multiple comparison). **D**, HCT116 cells were allowed to migrate on a collagen-coated rigidity gradient (0.1-80 kPa), previously conditioned or not by CAFs and treated or not with Cilengitide, as indicated. HCT116 cells were imaged every 20 min for 14 hours. Cells were manually tracked and the forward migration index (FMI) was calculated using the Chemotaxis tool plugin in FIJI software. Values close to 0 (blue line) indicate a random migration. Orientation of the gradient is shown. A total of 514, 1039 and 465 HCT116 cells were tracked from at least three independent experiments. Data are expressed as mean FMI ± SD (ANOVA one way, Dunnett’s multiple comparison).

## DISCUSSION

Overall, our results show that migrating fibroblasts deposit membranous material in the form of tracks and that cancer cells can use these tracks to steer their migration. CCSs play an instrumental role in this process as they are required to adhere to tracks. Collagen fibers and tracks share common traits such as their tubular shape and diameter. The difference, however, is that CCSs cannot internalize collagen fibers because these structures are too long and too rigid to be accommodated into clathrin-coated pits (Elkhatib et al., 2017). However, in the case of tracks, which are membranous structures and are thus less rigid than collagen fibers, CCSs can internalize portions of them with the necessary assistance of actin dynamics. Thus, tracks may represent an intercellular communication system that is read and erased by CCSs of following cancer cells. In addition to the tracks described here, and migrasomes that originate from tracks, deposition of secreted extracellular vesicles has been proposed to draw paths for the migration of following cells (Sung et al., 2020). Also, paths made of fibronectin deposited by migrating cells were shown to represent self-induced environmental perturbations that influence future cell trajectories (d’Alessandro et al., 2021). Thus, migrating cells may assemble different types of paths that influence self and/or neighboring cell migration. We propose that tracks encode a long-lived but erasable spatial memory of CAF migration trajectories that instructs cancer cell migration.

## MATERIALS AND METHODS

### Cell lines and constructs

hTert-immortalized Cancer associated fibroblasts (CAFs) stably expressing mOrange-CAAX (a gift from D. Matic Vignjevic, Insitut Curie, Paris, France) and desmoplastic small round cell tumor’ CAFs (a gift from C. Hénon, Gustave Roussy) were grown in DMEM Glutamax supplemented with 10% foetal calf serum and 1% insulin-transferrin-selenium (ITS ref. I3146, Sigma) at 37°C in 5% CO2. Wild type and β1-integrin Knock Out immortalized mouse osteoblasts (a gift from C. Albiges-Rizo - IAB Grenoble, France), MDA-MB-231 cells (a gift from P. Chavrier, Institut Curie, Paris, France) and genome edited MDA-MB-231 cells engineered to expressed an endogenous GFP-or mCherry-tagged μ2 subunit (a gift from D. Drubin, University of California-Berkeley, California, USA), HEK293T (a gift from P. Chavrier, Institut Curie, Paris, France), and HCT116 (A gift from D. Matic Vignjevic, Institut Curie, Paris, France) were grown in DMEM Glutamax supplemented with 10% foetal calf serum at 37°C in 5% CO2. All cell lines were periodically tested for the absence of mycoplasma contamination. All experiments were performed on hTert-immortalized Cancer associated fibroblasts (CAFs) stably expressing mOrange-CAAX, unless otherwise stated.

Substrates were coated with 50 μg/ml collagen-I (Thermo Fisher Scientific - ref. A10483-01), 20 μg/ml fibronectin (Corning – ref. 354008), or left uncoated.

Plasmids encoding for β5-integrin were described previously (Baschieri et al., 2018). Briefly, PCR purified integrin β5 with engineered flanking restriction sites (Xho-I/BamHI) was subcloned into the multi-cloning sites of pEGFP-N1 (Clontech) to encode an in-frame fusion protein with the carboxy-terminal EGFP-tag (pEGFP-N1-Integrin β5) or with carboxy-terminal mCherry-tag (mCherry-N1-Integrin β5). Thy1.1-GFP was a gift from Britt Glaunsinger (Addgene plasmid #131703). Lact-C2-GFP (MFGE8) was a gift from Sergio Grinstein (Addgene plasmid #22852). mRFP-Lact-C2 was a gift from Sergio Grinstein (Addgene plasmid #7406). pHR GFP-CAAX (for lentiviral production) was a gift from Ron Vale (Addgene plasmid #113020).

Lentiviral plasmids encoding for Integrin β3-GFP, integrin β1-GFP (built as in Huet-Calderwood et al., 2017), integrin αv-mCherry, integrin β5-BFP and Vinculin-GFP were purchased from VectorBuilder and are available upon request.

Lentiviruses were produced in HEK293T cells using the packaging plasmid psPAX2 and the VSV-G envelope expressing plasmid pMD2.G according to the Trono lab protocol. To generate stable cell lines, cells were transduced with lentiviruses and sorted via fluorescence-activated cell sorting. Plasmids were transfected 24 h after cell plating using either Lipofectamine 3000 according to the manufacturer’s instructions (MDA-MB-231) or by electroporating cells in suspension using AMAXA nucleofector Kit V program A-024 (CAFs) or program X-001 (osteoblasts) according to the manufacturer’s instructions. Alternatively, linear PEI (MW 25.000 – Polysciences Cat. Nr. 23966) at 1 mg/ml was used to transfect 50 % confluent cells in a 6 well plate according to the following protocol: 2 μg of DNA were added to 100 μl of OptiMEM, followed by addition of 4 μl of PEI, vortex and incubation for 10 minutes at RT prior to add the mix to the cells.

### Antibodies and drugs

Rabbit polyclonal anti-α-adaptin antibody (ref. sc-10761) was purchased from Santa Cruz Biotechnology Inc. (Santa Cruz, CA, USA). Mouse monoclonal anti-α-adaptin antibody (ref. ab 2807), rabbit polyclonal anti-Talin1 (ref. ab71333), and mouse monoclonal anti-integrin αvβ3 (ref. ab190147) were purchased from Abcam. Mouse monoclonal anti-clathrin heavy chain (CHC – ref. 610500) antibody and mouse monoclonal anti-integrin αv (ref. 611012) were obtained from BD Transduction Laboratories (Becton Dickinson France SAS, Le Pont-De-Claix, France). Mouse monoclonal anti-α-Tubulin (ref. T6199) was purchased from Merck. Rabbit anti-Integrin β1 was a gift from the Albiges-Rizo’s lab. Rabbit monoclonal anti-Integrin β5 (ref. 3629) was purchased from Cell Signaling Technology. Mouse monoclonal anti-integrin αvβ5 (ref. MAB1961) and integrin β3 (ref. AB2984) were purchased from Millipore. HRP-conjugated anti-mouse (ref. 115-035-062) for western blot was from Jackson ImmunoResearch Laboratories (West Grove, PA, USA). HRP-conjugated anti-rabbit (ref. A0545) was purchased from Sigma.

Alexa-conjugated secondary antibodies anti-mouse-A488 (ref. A21208), anti-mouse-A647 (ref. A31571), anti-rabbit-A647 (ref. A31573) and anti-rabbit-A488 (ref. A21206) were from Molecular Probes (Invitrogen). Alexa-conjugated antibodies A568 (ref. ab175471) was obtained from Abcam. Anti-mouse CF633 (ref. 20124) was from Biotium. Phalloidin A488 (ref. A 12379), Phalloidin-A633 (ref. A22284), WGA-A633 (ref. W-21404), WGA-A488 (ref. W-11261) were from Molecular Probes (Invitrogen). For STORM microscopy, CF647 F(ab’)2 (ref. BTM20042) and CF680 F(ab’)2 (ref. BTM20064) were bought from Ozyme.

Rat tail Collagen-I (ref. A10483-01) and Fibronectin (Corning - 354008) were purchased from GIBCO. Cytochalasin D (ref 22144-77-0) was purchased from Sigma and used at a final concentration of 10 μM. Cilengitide was purchased from Selleckchem (Cat. Nr. S7077) and used at a final concentration of 10 μM. Calyculin A (ref. PHZ1044) and Calcein AM (ref. 65-0853-39) were purchased from Thermofisher and used at a final concentration of 20 nM and 1 μM respectively. For Western Blot experiments, cells were lysed in ice cold MAPK buffer (100mM NaCl, 10 nM EDTA, 1% IGEPAL ® CA-630, 0.1% SDS, 50mM TRIS-HCl pH 7.4) supplemented with protease and PhosSTOP phosphatase inhibitor (ref. 04 906 845 001 – Roche). Antibodies were diluted at 1:1000 in PBS - 0.1% Tween - 5% non-fat dried milk.

### RNA interference

For siRNA-mediated depletion, 150 000 cells were plated together with the indicated siRNA (30 nM) using RNAimax (Invitrogen, Carlsbad, CA) according to the manufacturer’s instruction. Protein depletion was checked after 72 h of siRNA treatment by western blot or immunofluorescence with specific antibodies. To perform depletions of AP2, Clathrin heavy chain, and Talin1 in MDA-MB-231, as well as to perform depletions in CAFs, cells were transfected once as described above and then a second time, 48 hours later, with the same siRNAs. In this case, cells were analyzed 96 hours after the first transfection. The sequences of the siRNAs used are reported in Table 2.

### Indirect immunofluorescence microscopy and fluorescence quantification

Cells were fixed in ice-cold methanol or 4% PFA and processed for immunofluorescence microscopy by using the indicated antibodies. For anti-Talin1 staining, cells were briefly pre-extracted for 30 sec using PEM buffer (PIPES 80 mM, EGTA 5 mM, MgCl2 2 mM. pH 6.8, 1% TritonX100) prior to PFA fixation. Coverslips were mounted with Fluoromount-G (ref. 0100-20 – ref. 0100-01 SouthernBiotech).

Immunofluorescence images were acquired on a Leica Thunder epifluorescence microscope (Leica Microsystems Ltd., Wetzlar, Germany) through 63x (N.A. 1.40), 20x (HC N.A. 0.40 PH1), 10x (Fluotar N.A. 0.32 PH1) or 5x (PLAN 5/ N.A. 0.12 PHO) objectives. The microscope was equipped with LED5 lamp from Leica, a filter cube DFT51010, an external wheel EFW for DMi8, and sCMOS K5 camera. A PeCon chamber i8 was installed on the microscope to perform live imaging at 37°C with 5% CO2. The microscope was steered by the Leica dedicated LasX software® with Navigator allowing to generate mosaic stitched images.

To monitor the stability of tracks over time, cells were plated on 35 mm glass-bottom dishes with grids of 50 μm (ref. P35G-1.5-14-C-GRID - MatTek Corporation). Images were acquired 2 days after cells plating. With the help of the grids, the same regions were imaged 72h later and the number of tracks still present on the surface was quantified.

To quantify CCSs and FAs enrichment along tracks, 12.000 CAFs were plated on 18 mm diameter coverslips for 48 h, then 60.000 genome edited MDA-MB-231 cells wt, engineered to expressed an endogenous GFP-tagged μ2 subunit, or stably expressing vinculin-GFP were added for 35 minutes. Cells were then fixed with PFA and integrin β5 was stained to visualize tracks. Alternatively, staining against integrin β5 and α-adaptin subunit of AP-2 or Talin1 was performed. Tracks under MDA-MB-231 were manually segmented and the fluorescence intensity of CCSs or FAs markers in the region corresponding to tracks was quantified. The masks were then shifted by 10 pixel to calculate the background, non-specific GFP average fluorescence intensity. The ratio between the GFP signal under the tracks vs the GFP signal in the shifted region was used to quantify enrichment of CCSs or FA along tracks. All quantifications on immunofluorescence images were done with FIJI after background subtraction.

### Acrylamide gels of controlled stiffness

Coverslips or glass bottom dishes (ibidi - ref. 81218-200) were incubated with APTMS (3-aminopropyltrimethoxysilane) for 15 min at RT, then washed extensively with water and incubated for 30 min with Glutaraldehyde 0.5% in PBS and washed again with water. Acrylamide 40% and bis-acrylamide 2% were mixed (respectively 5% and 0.04% for 0.1 kPa gels, 7.5% and 0.06% for 5 kPa gels, 18% and 0.4% for 31.7 kPa gels, and 16% and 0.96% for 80 kPa) with PBS, APS 10% and TEMED. 9 μl of this solution were polymerized on the treated glass. Gels were washed with PBS, followed by a 30 min incubation with 300 μl 50mM Hepes pH 7.5+ 100 μl sulfo-sampah (1mg/ml in 50mM Hepes pH 7.5) + 100 μl EDC (10mg/ml in 50mM Hepes pH 7.5). Gels were subsequently cross-linked under UV light for 10 minutes, washed and incubated with 50 μg/ml collagen-I overnight at 37°C. Elasticity of the different gels was controlled by Atomic Force Microscopy as indicated in (Betz et al., 2011) The generation of 80 kPa gels was performed according to a previously published protocol (Cozzolino et al., 2016). Durotaxis gradients were obtained by placing 4 μl of 0.1 kPa solution next to 4 μl of 80 kPa solution on a treated glass-bottom dish. Fluorescent beads of 500 nm diameter (ref. F8812 - Invitrogen) were added to one of the two solutions to visualize the gradient. An 18 mm diameter glass coverslip was then quickly placed on top of the droplets, thus allowing for mixing of the two solutions and generation of a rigidity gradient. This method generates rigidity gradients where the stiffness increases linearly for 1-2 mm, as verified by AFM. Durotaxis gradients were coated with 50 μg/ml collagen I overnight at 37°C. Imaging was started 5h after cell plating and performed at the center of the gradient which was determined by looking at the fluorescent beads inside the gels. The number of cells plated on a gradient was 50 000 for CAFs and 100 000 for the other cell lines. One fluorescence image was acquired to visualize the fluorescent beads and identify the center of the gradient. Then, videos of 12-14 hours were recorded at a frequency of 1 image every 20 min with phase contrast. Cells were manually tracked using the “Manual tracking” plugin of FIJI. Forward Migration Indexes were obtained with the Ibidi “Chemotaxis and Migration Tool” in FIJI.

### Live cell spinning disk microscopy and quantifications

Cells were imaged on a Nikon Ti2 Eclipse (Nikon France SAS, Champigny sur Marne, France) inverted microscope equipped with a 60x NA 1.40 Oil objective WD 0.130, a sCMOS PRIME 95B camera (Photometrics, AZ, USA) and a dual output laser launch, which included 405, 488, 561 and 642 nm 30 mW lasers. The emission filters characteristics are as follows 452/45nm (Semrock Part# FF01-452/45); 470/24 (Chroma 348716); 525/50nm (Semrock Part# FF03-525/50); 545/40nm (Chroma 346986); 6986); (Semrock Part# FF01-609/54); 708/75nm (Semrock Part# FF01 708/75). The microscope was steered by Metamorph 7 software (MDS Analytical Technologies, Sunnyvale, CA, USA).

For CCS dynamics quantification along tracks, cells were imaged at 1 image every 5 sec for 5 min. Tracks were manually segmented and CCSs lifetime was measured on tracks vs outside tracks using the TrackMate plugin of FIJI. Tracks below 5 seconds of duration (detected on only 1 frame) were discarded. To quantify CCSs nucleation, the number of CCSs appearing during the 5 min long video was counted and normalized to the area. At least 1000 tracks from at least 5 cells per condition and per experiments were quantified in 3-5 independent experiments. Data are expressed as mean ± SD. Videos of 8-12 hours were recorded at a frequency of 1 image every 5 min. When needed, prior to image analysis, videos were realigned either manually or using the Template Matching plugin of FIJI.

### TIRF microscopy

For total internal reflection fluorescent microscopy (TIRF), MDA-MB-231 cells transfected with the indicated plasmids were imaged through a 100x 1.49 NA APO TIRF WD 0.13-0.20 oil objective lens on a Nikon Ti2 Eclipse (Nikon France SAS, Champigny sur Marne, France) inverted microscope equipped with a sCMOS PRIME 95B camera (Photometrics, AZ, USA) and a dual output laser launch, which included 405, 488, 561 and 642 nm 30 mW lasers, and driven by Metamorph 7 software (MDS Analytical Technologies, Sunnyvale, CA, USA). A motorized device (Piezo electrical XYZ stage from Nikon) driven by Metamorph allowed the accurate positioning of the illumination light for evanescent wave excitation. Videos of 8-12 hours were recorded at a frequency of 1 image every 5 min. When needed, prior to image analysis, videos were realigned either manually or using the Template Matching plugin of FIJI.

### Interference Reflection Microscopy

Images were acquired on a Leica Sp8 confocal microscope with a Pecon incubation chamber equipped with two hybrid and three PMT detectors, using 405, 488, 561 and 633 nm lasers and an 85/15 cube. Imaging was performed using a 63× objective (1.4 NA) and the 633 nm laser was used to have the IRM picture. Fast Fourier Transforms (FFTs) of the raw images were obtained in FIJI and used to remove linear interferences. Final images were obtained using the inverse FFT command on FIJI.

### 3D migration in collagen networks

For 3D cell migration assays, a glass-bottom dish was treated with Poly-L-Lysine 0.1% (Sigma – Ref. P8920) for 5 min at RT prior to add a 50 μl droplet of collagen type I (BD bioscience - 50:1 ratio unlabeled vs Alexa548-labeled) at a final concentration of 2.2 mg/ml mixed with 5,000 CAFs CAAX-mOrange cells. Collagen was allowed to polymerize at room temperature for 30 min before to immerge the setup in pre-warmed DMEM supplemented with FCS and ITS. 24h later, cells were imaged by spinning disk microscopy.

### Proteomic analysis of CAF-tracks

CAFs were plated at a density of 40 cells/mm^2^ in six T75 cell culture flasks. After 48 hours of migration, DMEM + ITS + FCS was replaced with DMEM + ITS. 24 h later, Calyculin A was added (final concentration 20 nM) for 20 minutes to induce cell detachment from the substrate. Flasks were then washed 5 times with PBS. 3ml of DMEM were added to each flask. Cilengitide, a selective inhibitor of the integrins αvβ3 and αvβ5, was added to three flasks to harvest tracks (final concentration 20 μM), while DMSO was added to the other three flasks. After 80 min incubation at 37°C, the medium was recovered and centrifuged at 20.000g for 30 min. Pellets were then recovered in 20 μl of PBS and subjected to further analysis.

Each sample was dried and solubilized in 10 μL of 8M urea, 200 mM ammonium bicarbonate and then reduced in 5 mM dithiothreitol, pH 8 with vortexing at 57°C for 30min. After cooling to room temperature, cysteines were alkylated by adding 10 mM of iodoacetamide for 30 min in the dark. After diluting to 1 M urea with 200 mM ammonium bicarbonate pH 8.0, samples were digested with 0.4μg trypsine/LysC (Promega) overnight, with vortexing at 37°. Samples were then loaded onto homemade C18 StageTips for desalting. Peptides were eluted using 40/60 MeCN/H2O + 0.1% formic acid, vacuum concentrated to dryness and reconstituted in 10μl or 20μl injection buffer (0.3% TFA) before nano-LC-MS/MS analysis. Online chromatography was performed using an RSLCnano system (Ultimate 3000, Thermo Fisher Scientific) coupled to an Orbitrap Fusion Tribrid mass spectrometer (Thermo Fisher Scientific). Peptides were trapped on a C18 column (75 μm inner diameter × 2 cm; nanoViper Acclaim PepMapTM 100, Thermo Fisher Scientific) with buffer A (2:98 MeCN:H2O in 0.1% formic acid) at a flow rate of 3.0 μl/min over 4 min. Separation was performed on a 50 cm x 75 μm C18 column (nanoViper Acclaim PepMapTM RSLC, 2 μm, 100Å, Thermo Scientific), regulated to a temperature of 40 °C with a linear gradient of 3% to 32% buffer B (100% MeCN in 0.1% formic acid) at a flow rate of 150 nl/min over 91 min. Full-scan MS was acquired using an Orbitrap Analyzer with the resolution set to 120,000, and ions from each full scan were higher-energy C-trap dissociation (HCD) fragmented and analysed in the linear ion trap. For identification, the data were searched against the Homo Sapiens UP000005640 database using Sequest HT through Proteome Discoverer (v.2.4). Enzyme specificity was set to trypsin and a maximum of two miss cleavages sites were allowed. Oxidized methionine, Met-loss, Met-loss-Acetyl and N-terminal acetylation were set as variable modifications. Carbamidomethylation of cysteins were set as fixed modification. Maximum allowed mass deviation was set to 10 ppm for monoisotopic precursor ions and 0.6 Da for MS/MS peaks. False-discovery rate (FDR) was calculated using Percolator (The et al., 2016) and was set to 1% at the peptide level for the whole study. The resulting files were further processed using myProMS (Poullet et al., 2007) v.3.9.3 (https://github.com/bioinfo-pf-curie/myproms). Protein were considered track candidates only if i) never found in PBS samples and ii) present in both track replicate with at least 3 peptides in each analysis. The mass spectrometry proteomics data have been deposited to the ProteomeXchange Consortium via the PRIDE (Perez-Riverol et al., 2019) partner repository with the dataset identifier PXD034091.

### Super resolution microscopy 3D STORM

For 3D STORM microscopy, immunolabelled cells were prepared on 1.5H coverslip using freshly prepared STORM buffer (Smart Kit, Abbelight, Cachan) and sealed with a silicon gasket. The sample was imaged through a 100x 1.49 NA APO TIRF WD 0.13-0.20 oil objective lens on a Nikon Ti2 Eclipse (Nikon France SAS, Champigny sur Marne, France) inverted microscope associated to a SAFe 360 detection module (Abbelight, Cachan) combined to two sCMOS Flash 4 v3 camera (Hamamatsu, Japan). Uniform and large field of view TIRF excitation is provided through an ASTER module (Mau et al., 2021), which included 405 nm (LBX-405-50-CSB-PPA, Oxxius 50 mW) and 640 nm (ERROL laser 500 mW) lasers and a quad band dichroic/emission filter (Semrock 405/488/532/640 refs. FF01-446/510/581/703-25 and Di03-R405/488/532/635-t1-25×36). Acquisition is driven by Neo software (Abbeligth, Cachan) and typically between 20000 and 40000 images (20 ms acquisition time) were acquired to reconstruct the final 3D super-resolved images. Two 3D imaging modalities were used. For single protein observation, SAFe detection module was configured for DONALD microscopy (Bourg et al., 2015), i.e. the fluorescence image is divided in two imaging path on the two camera by a 50-50% beamsplitter, and intrinsic supercritical angle fluorescence (SAF) is used to extract absolute axial position of single molecule events. For simultaneous dual protein observations labelled with CF647 and CF680, the SAFe module was configured for spectral demixing (Lampe et al., 2012), i.e. the fluorescence image is divided in two complementary spectral images by a dichroic with a cut-off wavelength at 700 nm (Chroma T700lpxr-3), and axial information is provided by two astigmatic lens in each imaging path (Cabriel et al., 2019). 3D STORM analysis including drift correction, axial information calculation (SAF or astigmatism), ratiometric measurements for spectral demixing of CF647 and CF680, filtering by cluster analysis were performed with Neo analysis software (Abbelight Cachan). Chimerax was repurposed for 3D surface rendering of 3Ddata in Movie 3 (Pettersen et al., 2021).

### Atomic Force Microscopy

Spectroscopy force experiments were performed in water with an Icon AFM coupled to Nanoscope V controller from Bruker. The probe was an OTR8 (Bruker) with a spring constant of 0.61N.m. The system was calibrated according to the manufacturer’s procedure: spring constant, deflection sensitivity and tip size. The approach/retract curves were acquired with a maximum load of 80 N and at a frequency of 1Hz. The approach/retract curves are processed using software Analysis Nanoscope (Vers. 2.0 Bruker). The moduli represent the average of 4 values calculated with the Sneddon model on retract curves.

### Statistical analyses

Graphics were prepared with the open-source software Instant Clue (Nolte et al., 2018) and Prism V8.0 and statistical analyses were performed using Prism V8.0 or Instant Clue software.

## ACKOWLEDGEMENTS

We thank the imaging facilities of Gustave Roussy for help with image acquisition and the microscopy and analysis platform of CYU Cergy Paris Université for help with AFM acquisition. We thank the imaging facilities of Gustave Roussy for help with image acquisition. Core funding for this work was provided by the Gustave Roussy Institute and the Inserm and additional support was provided by grants from the Agence Nationale de la Recherche (ANR-MOTILGUT), from Institut National du Cancer (2018-1-PL BIO-02-IGR-1) and from Fondation pour la Recherche Medical (DEQ20180339205) to GM. FB. was supported by the 2020 fellowship from Fondation Tourre and acknowledges support by the 2019 Metastasis Award from the Beug Foundation and by “dotation nouveau recruté 2022” from INSERM.

