## Supplementary information and figures for "Fibroblasts generate topographical cues that steer cancer cell migration"

### Supplementary Figure Legends

**Supplementary Figure 1. Characterization of CAF-tracks. A**, CAAX-mOrange-expressing CAFs (Blue) were embedded in a 3D network composed of collagen fibers (Red) and allowed to migrate for 24 hours. Cells were then imaged by spinning disk microscopy. Scale bar: 20 µm. **B**, CAAX-mOrange-expressing CAFs were allowed to deposit tracks on glass and tracks were imaged by spinning disk microscopy upon CAFs disconnecting from the tracks. Scale bar: 10 µm. **C**, Super resolution microscopy analyses of β5-integrin-BFP tracks. Scale bars, x-y plane: 10 µm, x-z plane: 100 nm. A color-coded scale indicates the distance from the glass in nm. **D**, Average track tubules diameter in x-y and x-z planes as measured by super resolution microscopy as in C. Data are expressed as mean diameter ± SD.

**Supplementary Figure 2. Characterization of tracks integrins content. A**, **B**, CAFs migrating on glass were fixed and stained for β5-integrin (blue) and phalloidin (A, red) or tubulin (B, red). Scale bars: 10 µm. **C**. CAFs migrating on glass were incubated with Calcein AM and imaged by confocal microscopy (left panel) and Interference Reflection Microscopy (right panel). Scale bar: 10 µm. **D**, Western-blot analysis of β5-integrin expression in CAFs treated with the indicated siRNAs as described in the Materials and Methods. Tubulin was used as a loading control. Molecular weights are indicated. **E**, Representative immunofluorescence images of CAFs treated with the indicated siRNAs and stained for αvβ5 integrin. Scale bar: 30 µm. **F**, Representative immunofluorescence image of β1-integrin-depleted CAF marked with Alexa-488-labelled Wheat Germ agglutinin. Scale bar: 20 µm. **G**, Western blot analysis of β1-integrin expression in CAFs treated with the indicated siRNAs. Tubulin was used as a loading control. Molecular weights are indicated. **H**, Representative immunofluorescence image of β1-integrin-knockout osteoblasts marked with Alexa-488-labelled Wheat Germ agglutinin. Scale bar: 20 µm. I. Western-blot analysis of β1-integrin expression in wild type (WT) or integrin β1$^{-/-}$ mouse osteoblasts. Tubulin was used as a loading control. Molecular weights are indicated. **J**, Representative images of integrin β5-GFP staining in mouse osteoblasts (top left panel) or of αvβ5 integrin staining in the other indicated cell lines. Scale bars: 20 µm.

**Supplementary Figure 3. Analyses of focal adhesions and CCSs interactions with tracks. A**, MDA-MB-231 cells were allowed to spread on CAF-tracks for 35 min before to be fixed and stained for αvβ5-integrin and Talin1. Scale bar: 10 µm. **B**, Quantification of Talin1-associated fluorescence in track areas versus other areas of the plasma membrane in cells as in A. Quantifications were performed with aligned Talin1 and αvβ5-integrin signals (on tracks) as well as upon shifting the αvβ5-integrin signal by 10 pixels (10 pixels shift). 47 cells from 3 independent experiments were analyzed. A value of 1 (blue line) means no enrichment on tracks. Results are represented as mean ratio of track-associated versus non-track-associated Talin1 signal ± SD (Student's T-test). **C**, MDA-MB-231 cells transfected with the indicated siRNA were fixed and stained for Talin1. Scale bars: 20 µm. **D**, HCT116 cells were allowed to spread on CAF-tracks for 1 h before to be fixed and stained for β5-integrin (blue) and α-adaptin (red). Scale bar: 10 µm. **E**, Quantification of α-adaptin-associated fluorescence in track areas versus other areas of the plasma membrane in cells as in D. Quantifications were performed with aligned α-adaptin and β5-integrin signals (on tracks) as well as upon shifting the β5-integrin signal by 10 pixels (10 pixels shift). 47 HCT116 cells from 3 independent experiments were analyzed. A value of 1 (blue line) means no enrichment on tracks. Results are represented as mean ratio of track-associated versus non-track-associated α-adaptin signal ± SD (Student's T-test). **F**, MDA-MB-231 cells genome-edited to express µ2-adaptin-mCherry were allowed to adhere on tracks produced by mouse osteoblasts stably expressing β5-integrin-BFP for 30 minutes. Scale bar: 10 µm. **G**, Quantification of µ2-adaptin-mCherry associated fluorescence in track areas versus other areas of the plasma membrane in cells as in F. Quantifications were performed with aligned µ2-adaptin and β5-integrin signals (on tracks) as well as upon shifting the β5-integrin signal by 10 pixels (10 pixels shift). 29 cells from 3 independent experiments were analyzed. A value of 1 (blue line) means no enrichment on tracks. Results are represented as mean ratio of track-associated versus non-track-associated α-adaptin signal ± SD (Student's T-test). **H**, **I**, MDA-MB-231 genome-edited to express µ2-adaptin-mCherry were allowed to spread on β5-integrin-BFP osteoblast tracks and imaged every 5 sec for 5 min. Lifetime (H) and nucleation rates (I) of CCSs on tracks vs outside tracks was calculated. 17 cells from three independent experiments were analyzed. Results are expressed as mean ± SD (Student's T-test).

**Supplementary Figure 4. Analyses of mechanism of CCSs recruitment on tracks. A**, MDA-MB-231 cells genome-edited to express µ2-adaptin-mCherry (red) were allowed to spread on β5-integrin-BFP CAF-tracks (blue) in the presence or not of Cytochalasin D, as indicated, and imaged every 5 min for 3 hours by spinning disk microscopy. A still frame acquired after 30 min of spreading is shown. Scale bars: 10 µm. **B**, Kymographs corresponding to boxed areas in A. **C**, MDA-MB-231 treated or not with Cytochalasin D were allowed to spread on β5-integrin-BFP tracks and imaged every 10 min for 3 hours. Data represent the evolution over time of the mean β5-integrin-BFP fluorescence intensity ± SD from three independent experiments (Unpaired T test with Welch's correction – $p<0.0001$ as compared to Control). **D**, Western-blot analysis of β1- and β5-integrins expression in CAFs cell lysate versus enriched CAF-tracks fraction. Tubulin was used as a loading control. Molecular weights are indicated. **E**, Quantification of μ2-adaptin-mCherry associated fluorescence in track areas versus other areas of the plasma membrane in MDA-MB-231 cells transfected with the indicated siRNAs. At least 15 cells per condition were analyzed from 3 independent experiments. A value of 1 (blue line) means no enrichment on tracks. Results are expressed as mean ratio of track-associated versus non-track-associated μ2-adaptin signal ± SD (uncorrected Fisher's LSD test). **F**, CAF stably expressing CAAX-mOrange and overexpressing MFGE8-GFP or Thy1-GFP were allowed to migrate on glass and imaged by spinning disk microscopy. Scale bars 10 μm. **G**, Western-blot analysis of αv- and β3-integrins expression in MDA-MB-231 and HCT116 cells. Tubulin was used as loading control. Molecular weights are indicated. **H**, Western-blot analysis of αv-integrin expression in MDA-MB-231 treated with the indicated siRNAs. Tubulin was used as a loading control. Molecular weights are indicated. **I**, MDA-MB-231 genome-edited to express μ2-adaptin-mCherry (red) and stably expressing β3-integrin-GFP (yellow) were allowed to spread on β5-integrin-BFP tracks (cyan) in the presence of 10 μM Cytochalasin D and imaged every 5 min for 12 hours. Still picture at 30 min of spreading is shown. Scale bar 10 μm. **J**, Kymograph corresponding to the boxed area in I.

**Supplementary Figure 5. Analyses of rigidity gradient gels and durotaxis capacities. A**, Rigidity-gradient gels were prepared as described in the Materials and Methods. Relative measurements of rigidity were performed by Atomic Force Microscopy (AFM) moving orthogonally to the gradient from one extremity of the gel to the other. Results are expressed as mean ± SD. **B**, Image of a 0.1-80 kPa rigidity gradient with fluorescent beads embedded in the stiff part of the gel. Scale bar: 200 μm. **C**, MDA-MB-231 cells were allowed to migrate on a collagen-coated rigidity gradient (0.1-80 kPa), previously conditioned or not by osteoblasts, as indicated. MDA-MB-231 cells were imaged every 20 min for 14 hours. Cells were manually tracked and the forward migration index (FMI) was calculated using the Chemotaxis tool plugin in FIJI software. Values close to 0 (blue line) indicate a random migration. Orientation of the gradient is shown. 350 cells and 541 cells were tracked from three independent experiments, for the non-conditioned and osteoblast-conditioned conditions, respectively. Data are expressed as mean FMI ± SD (Unpaired Student's T-test).

### Supplementary Movies

**Movie 1. CAFs deposit tracks as they migrate.** CAF stably expressing the plasma membrane marker CAAX-mOrange was allowed to migrate on glass and imaged by spinning disk microscopy every 30 min for 12 hours. Scale bar: 10 µm.

**Movie 2. MDA-MB-231 cells migrate along tracks.** MDA-MB-231 cell stably expressing Vinculin-GFP and migrating along CAAX-mOrange-marked tracks and imaged by spinning disk microscopy every 5 min for 9 hours. Scale bar: 10 µm.

**Movie 3. CCSs of cancer cells wrap around CAF-tracks.** 3D rendering of super resolution microscopy analyzes of CCSs of MDA-MB-231 cells (marked with α-adaptin) aligning along and wrapping around a CAF-track (marked with αvβ5 integrin). Scale bar 500 nm.

**Movie 4. Actin dynamics is required for track uptake.** MDA-MB-231 genome-edited to express µ2-adaptin-mCherry (Red) were allowed to spread on β5-integrin-BFP tracks (Cyan) in the presence or not of 10 µM Cytochalasin D, as indicated. Images were acquired on a spinning disk microscope every 5 min for 2 hours and 45 min. Upper panels show merge images of µ2-adaptin-mCherry and β5-integrin-BFP channels. Lower panels show β5-integrin-BFP channel only. Scale bar: 10 µm.

**Movie 5. β3 integrin in MDA-MB-231 cells accumulate along tracks.** MDA-MB-231 genome-edited to express µ2-adaptin-mCherry (red) and stably expressing β3-integrin-GFP (yellow) were allowed to spread on β5-integrin-BFP tracks (cyan) and imaged by TIRF microscopy every 5 min for 2 hours and 45 min. Scale bar: 20 μm.

Supp. Figure 1

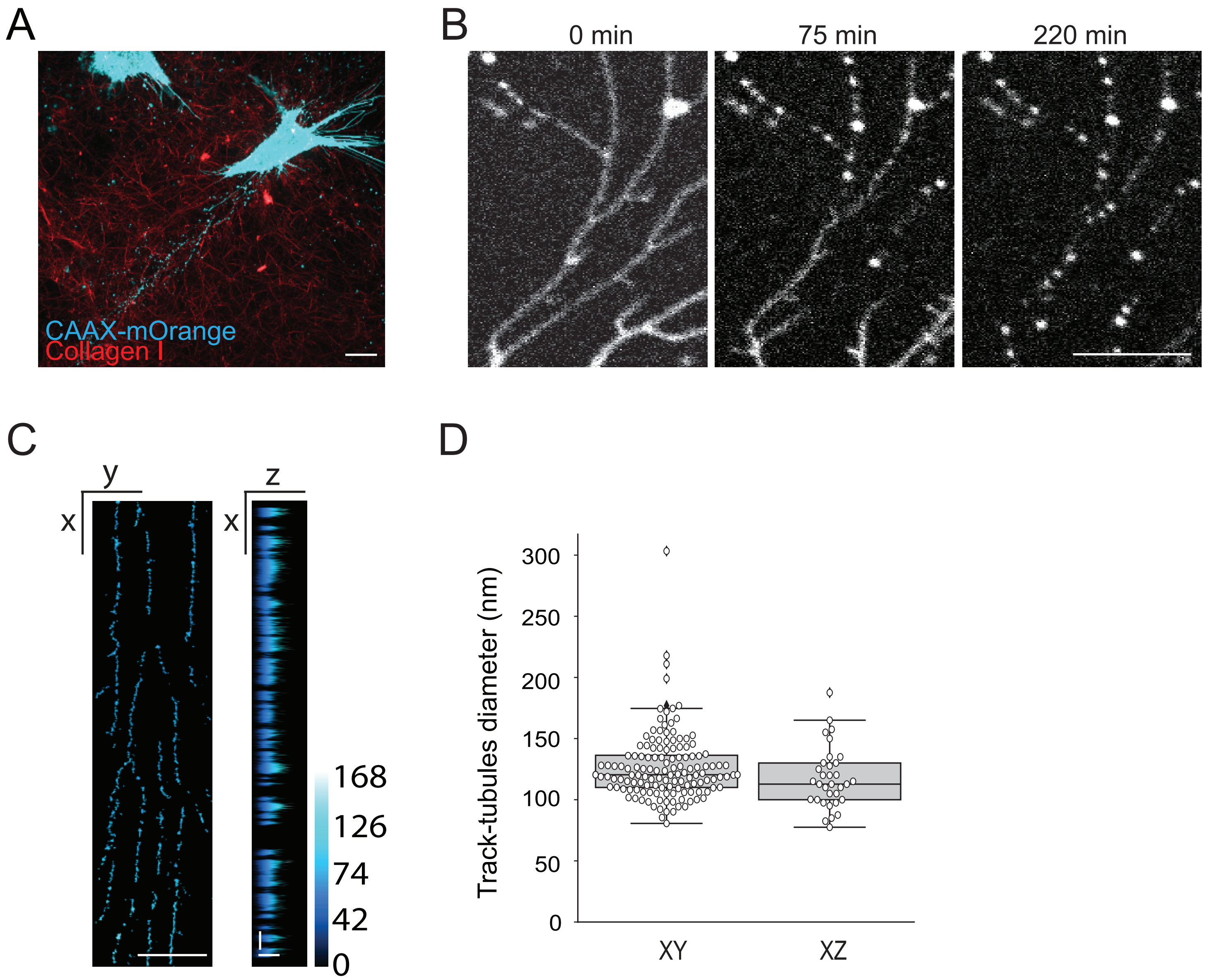

### Suppl. Figure 2

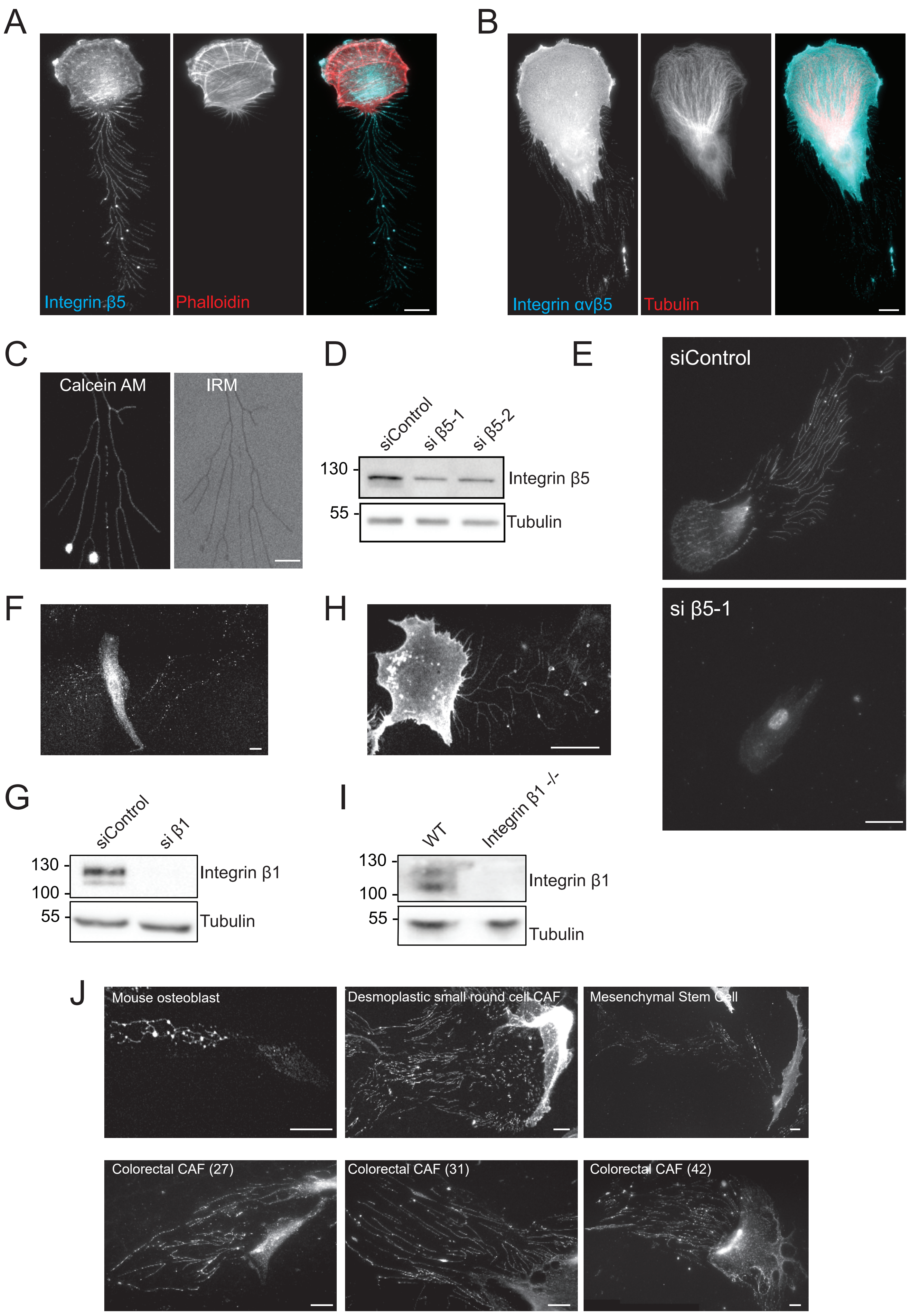

### Supp. Figure 3

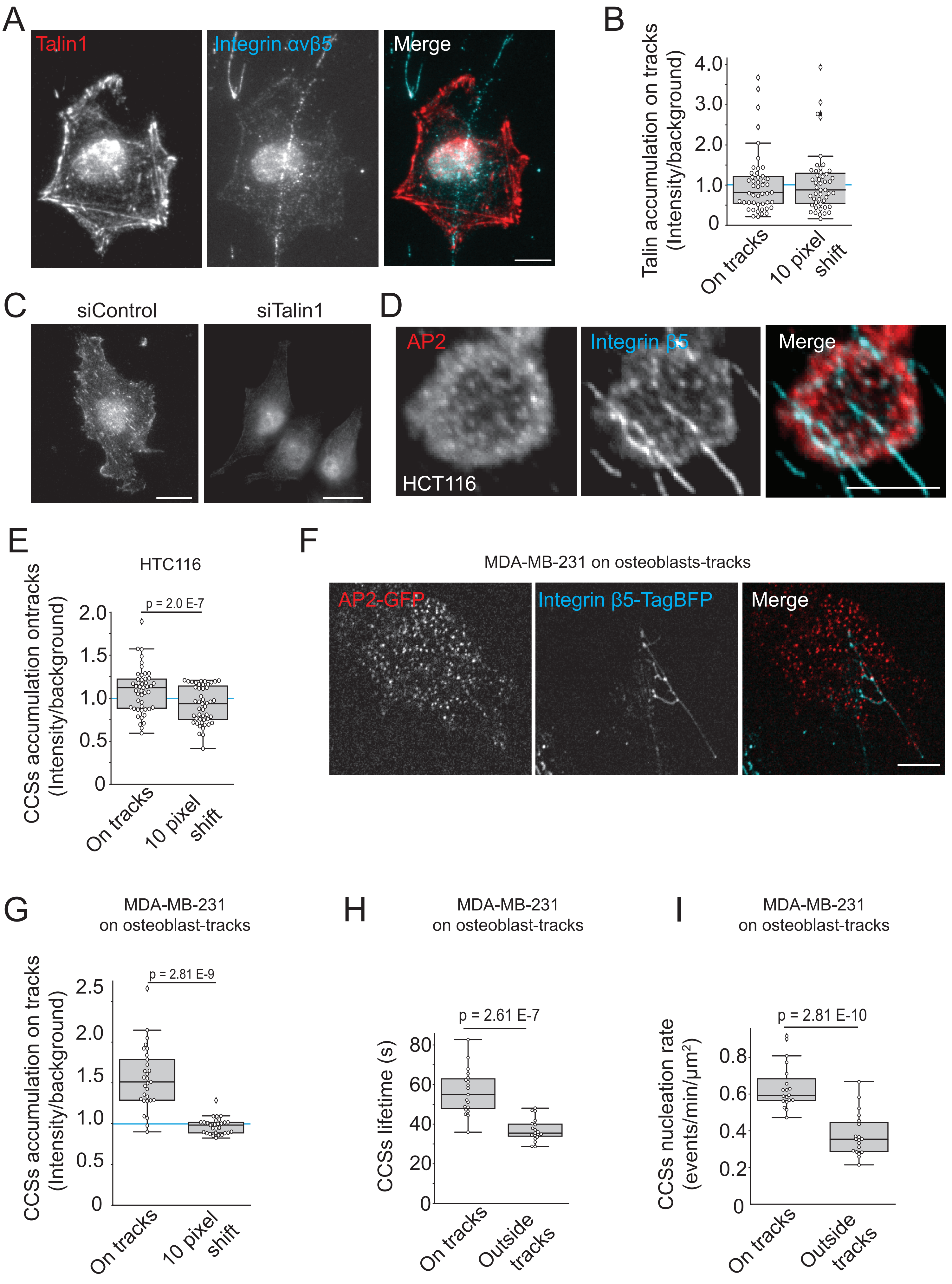

### Supp. Figure 4

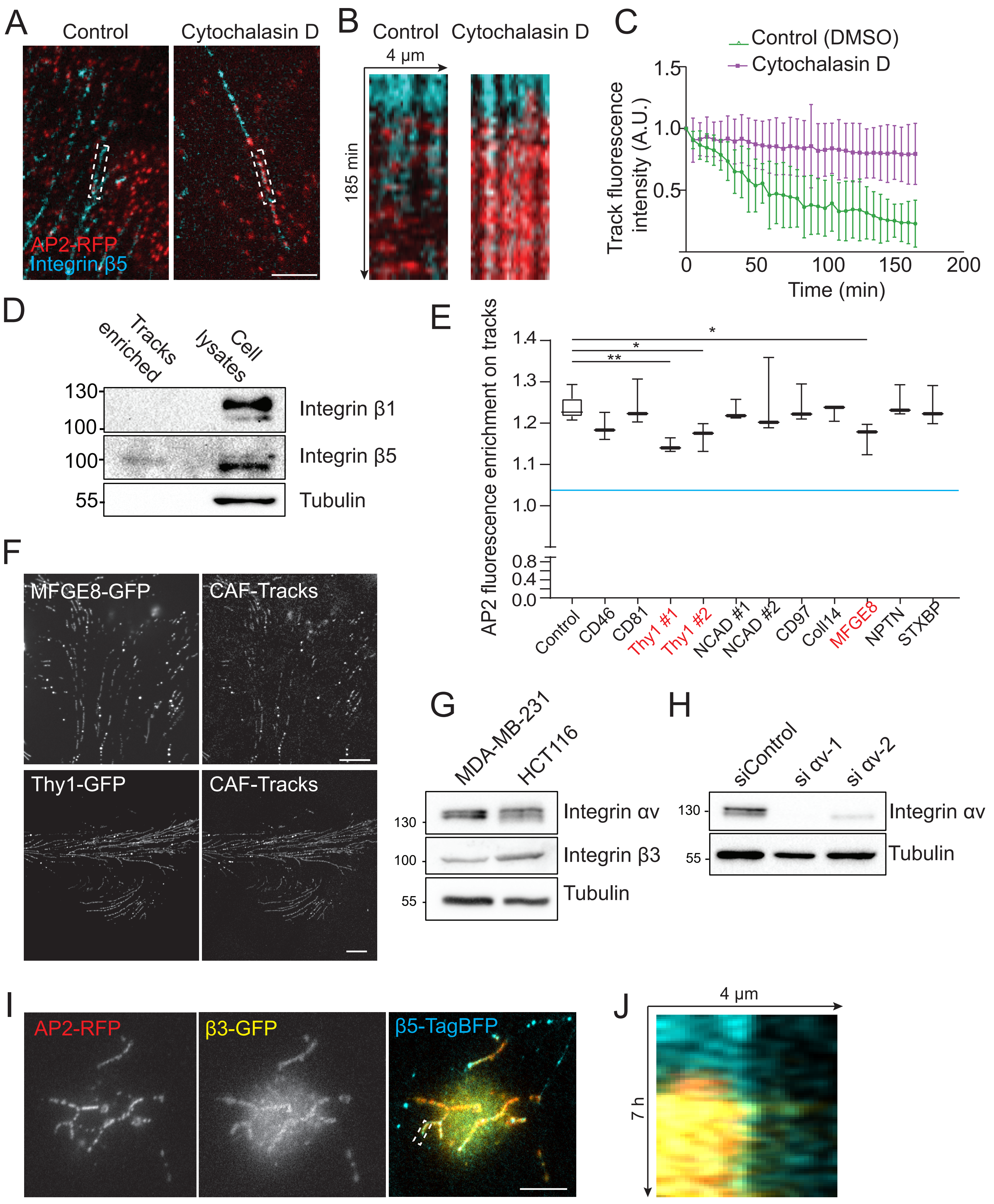

Supp. Figure 5

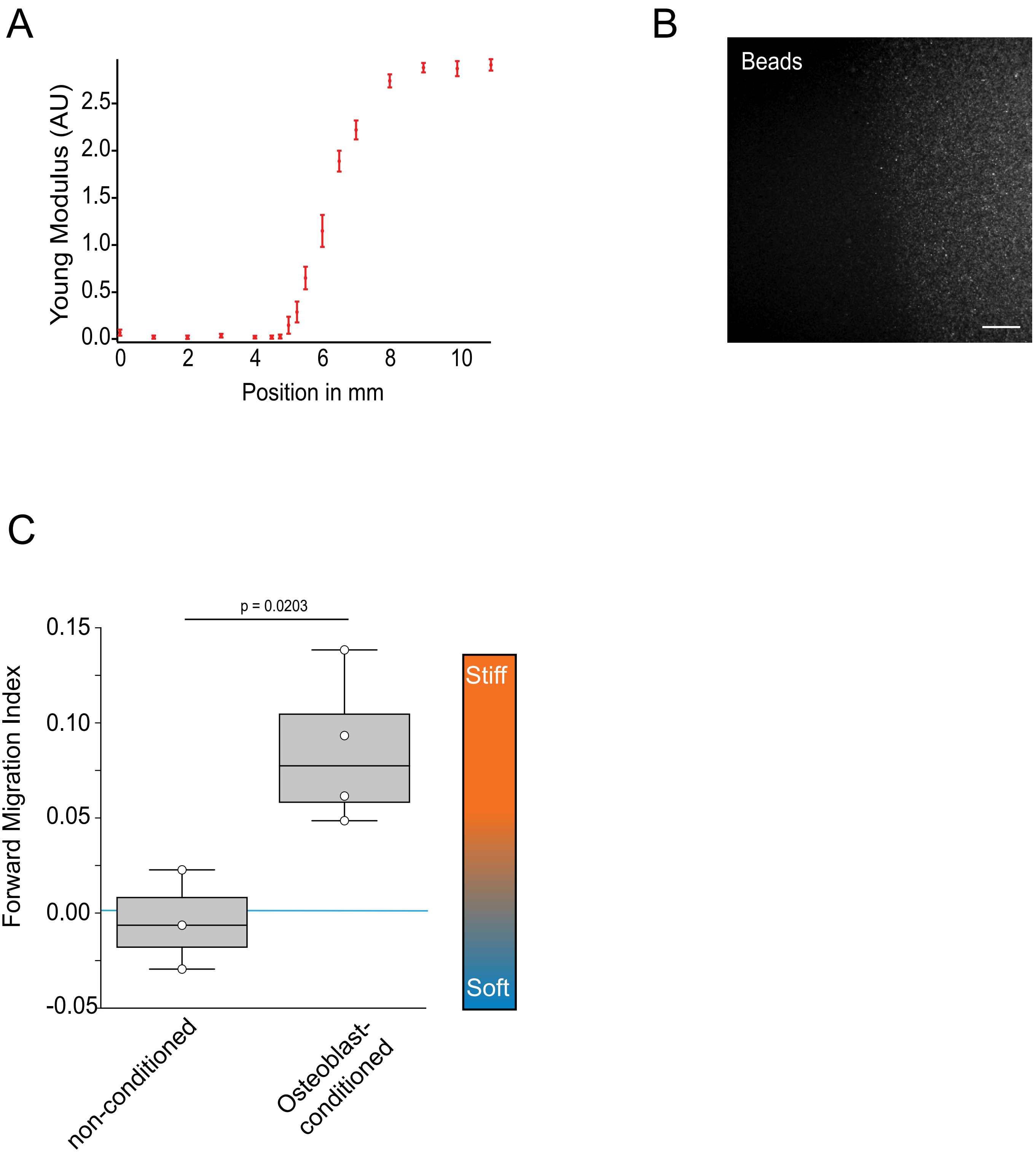
